# Host reverse transcriptase likely required to maintain persistent dengue virus infections in C6/36 insect cells

**DOI:** 10.1101/2022.12.20.521226

**Authors:** Warachin Gangnonngiw, Timothy W. Flegel, Nipaporn Kanthong

## Abstract

C6/36 cells challenged with Dengue virus serotype 2 (DENV-2 strain NGC) show initial cytopathic effects (CPE) they overcome within 3 split passages and resume normal growth despite persistent infection with DENV-2. We hypothesized that tolerated persistent infections also required persistent host reverse-transcriptase (HRT) activity, and this was subsequently proven using the RT inhibitor AZT (azidothymidine, a nucleoside analogue antiretroviral drug) to treat C6/36 cells challenged with DENV-1. We confirmed this using the RT inhibitor tenofovir disoproxil fumarate (TDF) with C6/36, U4.4cells challenged with DENV-1. However, it has never been determined whether the persistently tolerated infections are also dependent on persistent HRT activity. This brief report reveals that persistent HRT activity is required to maintain persistent DENV-2 infections. Specifically, TDF treatment (0.1 mM) revealed minor toxicity to naive C6/36 cells but led to cell death instead of accommodation upon concomitant challenge with DENV-2, as previously reported for DENV-1. However, TDF treatment of stable, grossly normal C636 cell cultures persistently infected with DENV-2 for up to 30 split passages reverted to CPE, supporting our earlier hypothesis that persistent host RT activity is also required to maintain persistent infections. This is a practical tool for research applications in virology of crustaceans and insects for screening living, grossly healthy, imported animals for listed viral pathogens.

## 1 INTRODUCTION

Recent research results have proven that insects and crustaceans are capable of a specific, adaptive immune response to viral pathogens based on nucleic acids instead of antibodies ^1,2^. These results support a proposal from 1998 that insects and crustaceans could tolerate one or more viruses in an adaptive response (termed viral accommodation) via unknown mechanisms ^3^. Viral accommodation was characterized by persistent infections with one or more pathogenic viruses without gross or histological signs of disease. These are not chronic or latent infections but instead continuous, persistent infections that may last for a host’s lifetime during which they are continuously infectious.

Previous research has shown that stable, grossly normal lines of persistently infected *Aedes albopictus* C6/ 36 mosquito cells can be produced in the laboratory by split-passage with the DNA virus *Aedes albopictus* densovirus (*Aal*DNV) and with the RNA viruses Dengue virus serotype 2 (DENV-2 strain NGC) and *Japanese encephalitis* virus (JEV). These may be single, dual or triple infections ^4,5^. However, the underlying mechanisms leading to these persistent infections and their maintenance in C6/36 cells are yet to be fully revealed.

We now know that host generated reverse transcriptase (RT) acting on invading viral RNA is a key enzyme in the viral accommodation response ^6,7^. It results in the production of highly variable fragments of viral copy DNA (vcDNA) in both linear (lvcDNA) and circular (cvcDNA) forms ^8^. These can give rise to small interfering RNA (siRNA) to feed both cellular and systemic antiviral responses based on RNA interference (RNAi) pathways ^1,2^.

Based on this recent progress on insect and crustacean antiviral immunity and on the importance of host generated RT in the process of establishing stable, persistent infections, we hypothesized that DENV-2 infections in the *Aedes albopictus* cell line would be dependent on host generated RT not only to establish tolerance to DENV-2 in persistent infections, but also to maintain them. Since this maintenance role had not yet been explored, we aimed to test the hypothesis by using the RT inhibitor tenofovir disproxil fumarate (TDF) to not only confirm that it would prevent the establishment of a stable *Aal-* DENV-2 infected cell line (similar to *Aal-*DENV-1) but that it would also destabilize a previously established, stable *Aal-*DENV-2 infected cell line, leading to its death.

TDF is in a class of medication called a nucleoside reverse transcriptase inhibitor (NRTI). It functions as a defective adenosine nucleotide that interferences with RT activity. The experiments described herein were possible with DEN-2 because it causes transient cytopathic effects in naive C6/ 36 cells that dissipate within 2 to 3 split passages after which the cells resume normal growth and morphology for unlimited passages despite 100% persistent infection with DENV-2 ^5^.

At the outset, we would like to emphasize that the sole and uncomplicated purpose of this short communication is to describe work proving that host RT-activity is required to not only induce tolerance to lethal viruses in insects but also to maintain tolerance.

## 2 RESULTS AND DISCUSSION

### 2.1 TDF had relatively low toxicity for C6/36 cells at 0.1 mM

Reverse transcriptase activity from endogenous retrotransposons is known in naïve C6/36 cells and is upregulated by RNA virus infection ^6^. After 24 hours of normal, naïve C6/36 cell treatment with TDF at the concentrations of 0.1, 0.25 and 0.5 mM, the number of viable cells decreased significantly (**Fig. 1**). The effect of TDF was not very severe at the lowest concentration tested (0.1 mM) that caused approximately 25% loss of relative viability by the MTT assay. Even treatment with 0.5 mM resulted in slightly less than 50% decrease in viability. This dose-related decrease in viability was associated with increasing numbers of small, dense, dying or dead cells with increasing concentrations of TDF (**Figs. 2B to 2D**). Thus, some cell death was significant for all treatments indicating that TDF itself had a negative effect on cell viability. It is important to notice that the cell death was not characterized by syncytial cell formation that is characteristic of death caused by DENV-2 at the early stages of infection prior to viral accommodation. Telomerase (a type of reverse transcriptase) is produced by C6/36 cells, and such inhibition has been related to induced mortality via apoptosis in cultured cancer cells ^11^. Thus, we may speculate that death (associated with the occurrence of small, dense, dead cells) was caused by TDF inhibition of host telomerase leading to apoptosis in naive C6/36 cells. Finally, because it was the least lethal, a 0.1 mM concentration of TDF was selected for use in tests on the effect of RT inhibition on the C6/36 cellular response to DENV-2.

**Figure 1.**
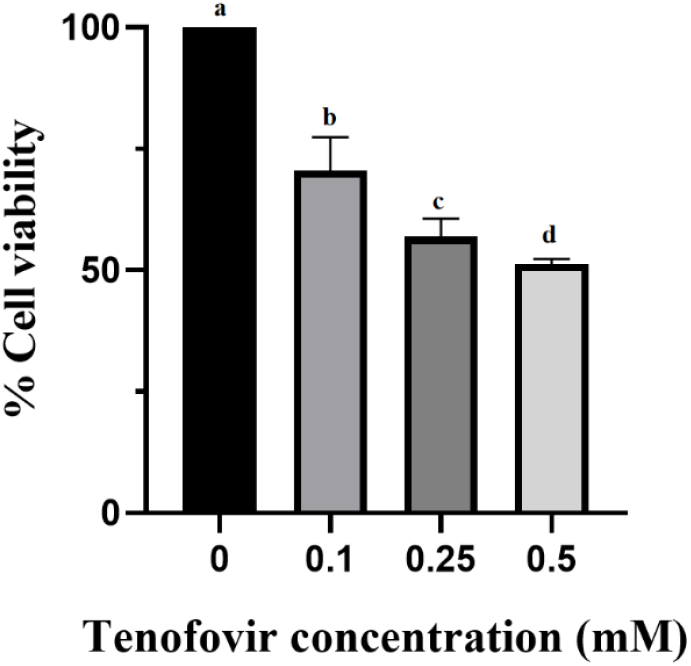
Bar graph of results from one-way ANOVA analysis of the MTT viability in C6/36 cell cultures treated or not with TDF for 24 hr. Baseline cell viability in the control wells not exposed to TDF was set at 100%. Data are expressed as percentage of the control. Bars with different letters are significantly different from one another (p<0. 05) and indicate increasing death with increasing concentrations of TDF.

**Figure 2.**
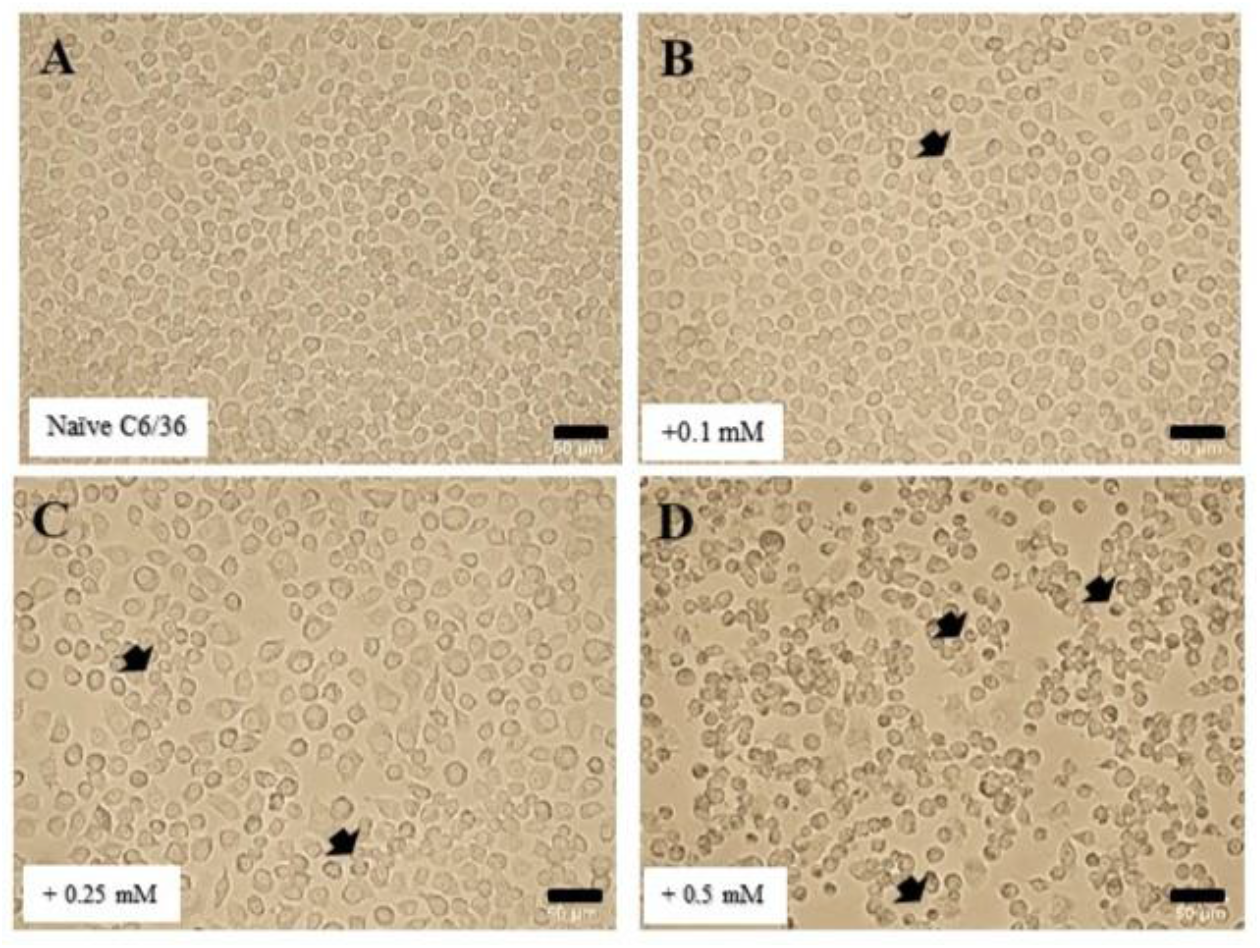
Naïve C6/36 cell morphology upon treatment or not with TDF for 24 h. **A.** Untreated naive cells. **B**. Cells treated with 0.1 mM and showing morphology similar to untreated cells except for the presence of a few smaller and dense, dead cells (black arrows). **C**. Cells treated with 0.25 mM and showing a mixture of normal cells like those in A and B in lower numbers but with more small dead cells (black arrows) than in A. **D**. Cells treated with 0.5 mM and showing the largest number of small dead cells. These photomicrographs correspond to the percentage of cell viability shown in Fig. 1.

### 2.2 RT inhibition by TDF prevents DENV-2 persistent infections

We wished to confirm that RT inhibition by TDF would have a similar effect to that described in earlier work using the RT inhibitor azidothymidine (AZT) in preventing persistent infections by DENV-1 in C6/36 cells ^6^. Thus, we tested whether TDF (a different inhibitor) would have the same effect on DENV-2 (a different virus). When monolayers of naive C6/36 cells were treated with TDF for 24 hours followed by DENV-2 challenge, cytopathic effects (CPE) were more severe in all the TDF treatment groups than in the untreated control. By passage 3, the number of cells had declined rapidly in the TDF treatment groups, and the severity of CPE increased (**Figs. 3A and 3B**). The untreated control cells showed minor cytopathic effects that corresponded to those previously reported and would be absent from the 4^th^ passage onward ^5^. Since all cells eventually died with DENV-2 challenge in the presence of TDF, persistently infected cultures with normal growth and morphology could not be obtained. Similar CPE-related phenomena were not described for the DENV-1 experiments with C6/36 cells previously reported ^7^.

**Figure 3.**
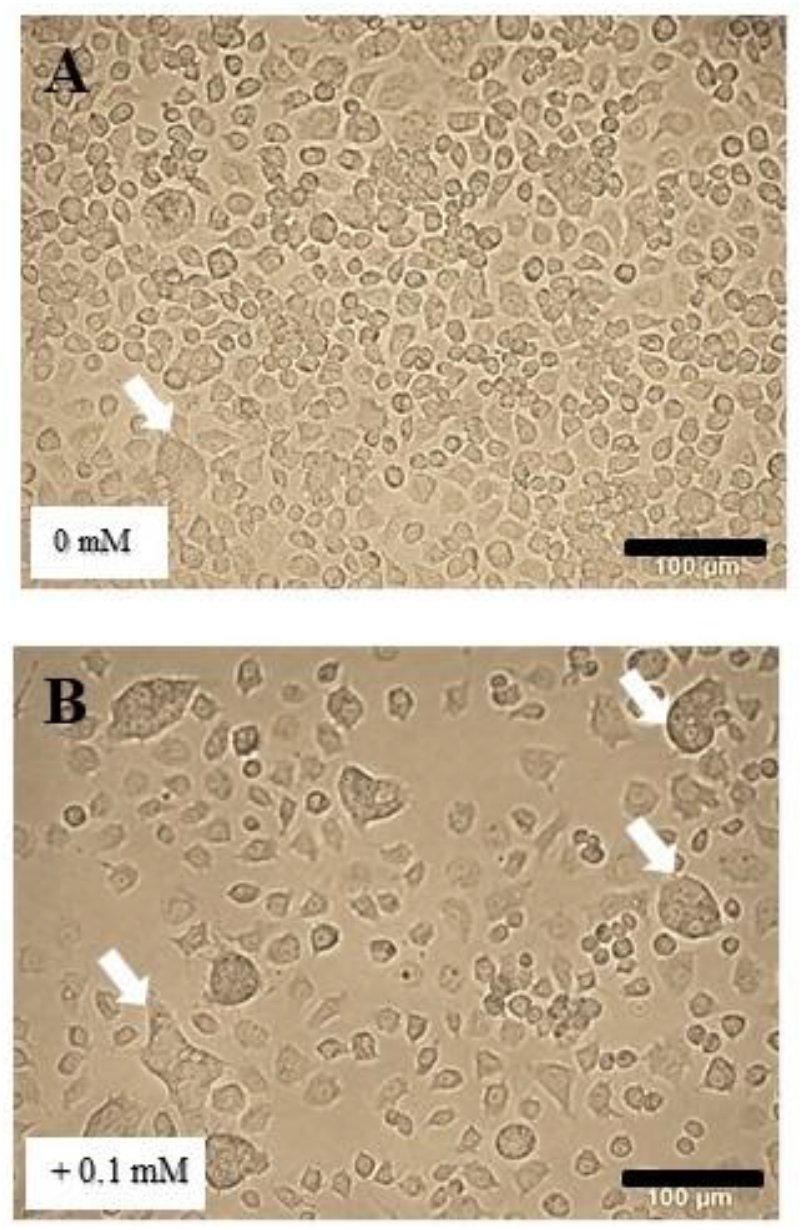
Naïve C6/36 cells morphology at Passage 3 in media containing TDF or not and challenged with DENV-2. **A.** DENV-2-challenged cells not exposed to TDF showing some minor CPE (white arrows) remaining at Passage 3, i. e., the cells are in the process of accommodating DENV-2. **B**. Treatment with 0. 1 mM TDF showing a decrease in cell numbers and an increase in CPE (white arrows) and the prevention of viral accommodation that occurred in the absence of TDF.

### 2.3 TDF increased DENV-2 production from naïve cells in a dose dependent manner

Measurement of DENV-2 copy numbers in the supernatant of C6/ 36 cell cultures at 48 hours after challenge with DENV-2 in the presence or absence of TDF revealed a significant increase in DENV-2 copy numbers dependent on the concentration of TDF (**Fig. 4**). The highest mean count was 9.60± 1.09 x10^6^ copies/µl with 0.5 mM TDF treatment while those in the untreated group and the 0.1 m TDF treatment were about 3.05± 0.43 x10^6^ copies/µl at 48 hours (Passage 0) after DENV-2 challenge. The increase in DENV-2 replication was positively related to the concentration of TDF. The results suggested that reduced cellular RT activity was associated with increased viral replication, as previously reported for DENV-1 in C6/36 cells ^7^. In contrast, in that report, the RNA virus Chikungunya did not respond to RT inhibition by increased viral replication. Thus, results may vary depending on the time of exposure before challenge and/or the concentration of RT inhibitor drug used in a specific host-viral strain interaction.

**Figure 4.**
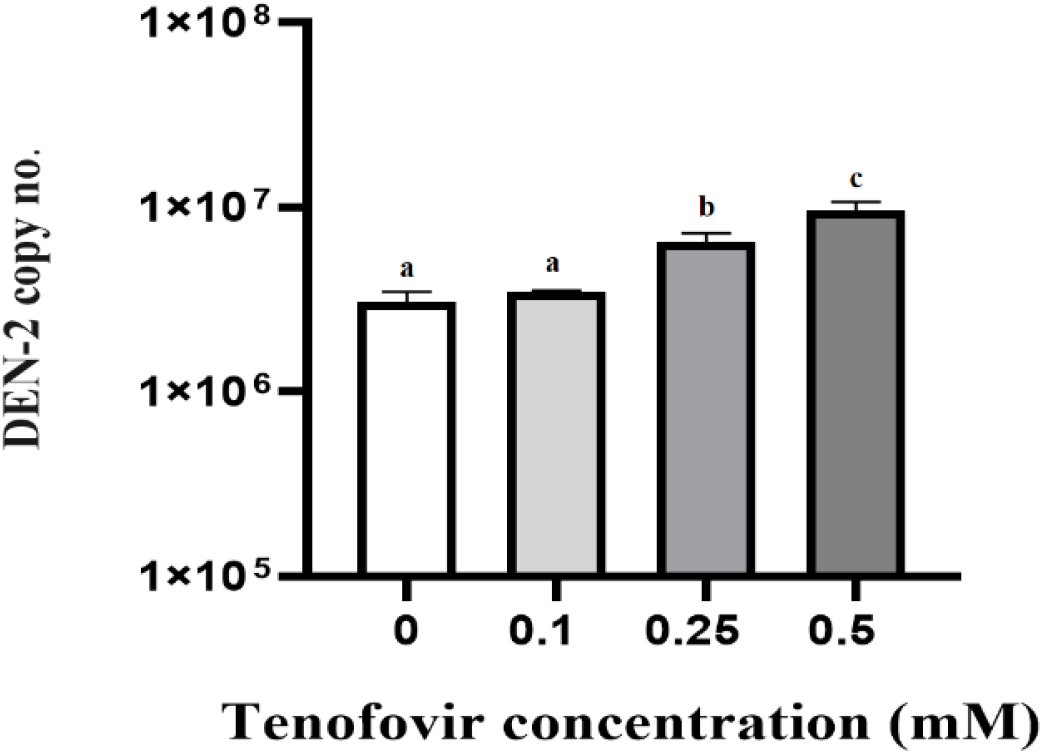
DENV-2 copy number in culture supernatant of naive C6/ 36 cells at 48 h in acute infection stage treated or not with TDF at different concentrations. Bars with different letters were significantly different (p≤0.05) from one another by one-way ANOVA.

### 2.4 TDF treatment destabilized persistently DENV-2-infected C6/36 cells

Our C6/36 cells persistently infected with DENV-2 by split-passaging 30 times and not treated with TDF (**Fig. 5A**) showed morphology like that of untreated, naive C6/ 36 cells as shown above in Fig. 2A. After 48 hours’ exposure to TDF, all cultures showed pathology with severity dependent on the TDF concentration. The grossly normal cells treated with 0. 1 mM TDF showed CPE comparable to that which occurs when untreated naive C6/36 cells are challenged with DENV-2 (see Fig. 3A above). It revealed that addition of TDF at 0. 1 mM (**Fig. 5B**) returned the persistently infected cells to a state resembling that of naive C6/36 exposed to DENV-2. It indicated that RT blockage destabilized viral tolerance, resulting in a return to CPE and cell death. Altogether, the results supported our hypothesis that that reverse transcriptase plays an essential role in maintaining, innocuous, persistent DENV-2 infections.

**Figure 5.**
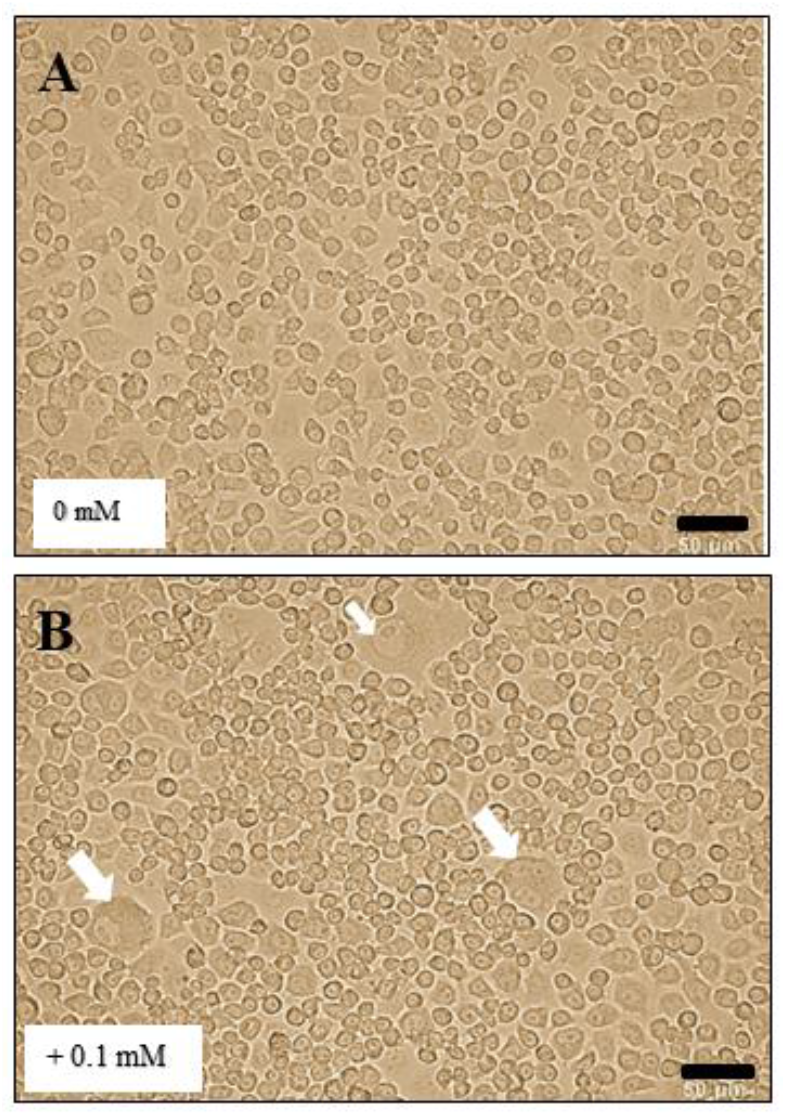

Moreover, DENV-2 replication increased in the presence of TDF. Although we did not purify a host reverse transcriptase from our C6/36 cells and carry out tests to prove inhibition by TDF, our results are consistent with previous published work revealing that RT inhibition drugs can prevent viral accommodation in cells of insects (Goic et al., 2013, 2016; Zhu et al., 2022), even though the detailed biomolecular mechanisms of the process have not yet been completely revealed. However, our results reveal some possible practical applications arising from our results in screening for accommodated viruses and increasing viral loads in experimental work. In addition the study may encourage more work on the detailed mechanisms of viral accommodation.

The DENV-2 copy number with 0.1 mM TDF treatment was three times higher (mean 7.64 ± 0.43 x10^6^ copies) than that in the untreated persistent infections (3.04± 0.43 x10^6^ copy) at the same passage (**Fig 6**). The formation of syncytia and the higher DENV-2 copy numbers with 0.1 mM TDF treatment revealed that inhibiting RT activity destabilized C6/36 cells tolerant to DENV-2. These results suggest that use of TDF treatment with persistently infected C6/ 36 cells at 0.1 mM TDF might be useful if high concentrations of DENV-2 were needed for some particular purpose.

**Figure 6.**
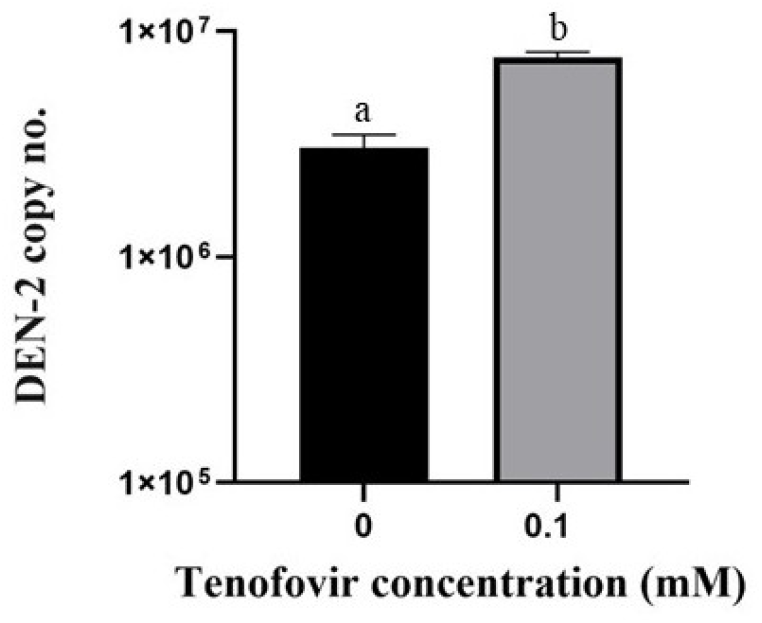
DENV-2 copy numbers in the culture supernatant of C6/ 36 cultures persistently infected with DENV-2 for 30 passages at 48h after being treated or not with TDF at 0. 1 mM. Bars with different letters were significantly different (p≤0.05) by paired t-test.

If it turns out that the requirement for RT activity is a general phenomenon upon which host insects and crustaceans depend to develop and maintain tolerance to viruses in persistent infections, it may have practical applications. For example, shrimp and insects often tolerate one or more lethal viruses that go unnoticed because they cause no signs of disease. However, those animals can transmit the viruses to naive individuals that may become infected and sometimes even die ^12^. This is very important for translocation of commercial insects (e. g., bees and silkworms) and crustaceans (e.g., shrimp) where simple quarantine cannot be used to reveal viral pathogens. In addition, tolerated viruses are sometimes present in very low numbers and may escape detection if very sensitive methods are not used. Thus, it is possible that treatment with an RT inhibitor during quarantine would result in disease expression or facilitate the detection of both known and unknown viral pathogens. This might be especially useful in the development of specific pathogen free (SPF) stocks derived from wild animals since signs of disease following TDF treatment might indicate the presence of even currently unknown viral pathogens.

## 3 CONCLUSIONS

Although it was previously known that host derived RT activity was necessary to establish stable, persistently infected cultures of insect cells as previously reported for DENV-1^13^, we have proven with DENV-2 that persistent RT activity is also required to maintain such stable cultures. Thus, we speculate that treating insects and crustaceans with an RT inhibitor may expose known or unknown, accommodated viruses in screening programs.

It may also be useful for increasing viral yields or increasing the severity of infections for research purposes. Recent studies have shown that host RT is also involved in production of viral copy DNA (vcDNA) from both RNA and DNA viruses in both linear and circular forms. The vcDNA, in turn, gives rise to small interfering RNA (siRNA) transcripts that lead to an RNAi response against both RNA viruses ^2^ and DNA viruses ^14,15^. In addition, the vcDNA also gives rise to host-cell genomic EVE, some of which produce negative-sense RNA that can also give rise to an RNAi response via PIWI proteins ^16,17^. Although C6/ 36 cells lack the gene Dicer-2 that is usually involved in the RNAi response, their ability to accommodate up to 3 viruses is unhindered ^4^. Thus, they constitute a good model for further studies on the detailed mechanisms underlying viral tolerance in the absence of Dicer-2.

## 3 MATERIALS AND METHODS

### 3.1 Statistical analysis

All experiments done out with the C6-36 cells were carried out in triplicate. Statistical analyses were performed using Graphpad prism v. 9 (GraphPad Software, San Diego, California USA, www.graphpad.com). For example, the quantity of viable cells and DENV-2 copy numbers were compared using one-way analysis of variance (one-way ANOVA) or paired t-tests, as appropriate. Differences with p-value ≤ 0.05 with 95% confidence were considered statistically significant. The one-way ANOVA results are provided in graphical form with quantitative bars marked with letters such that those with different letters are significantly different from one another while those with the same letter are not significant from one another.

### 3.2 Insect cell lines and DENV-2 inoculum

*Aedes albopictus* C6/36 cells (ATCC, catalogue number CRL-1660) and DENV-2 strain NGC were maintained and cultivated in the same manners as previously described ^9^. Briefly, C6/36 were grown in Leibovitz’s (L-15) medium containing 10% heat-inactivated fetal bovine serum (FBS), 10% tryptose phosphate broth (TPB) and 1.2% antibiotic (Penicillin G and Streptomycin). DENV-2 (NGC strain) stock was stored in 20% fetal bovine serum at −80°C. After thawing at room temperature, the stock was used as inoculum for monolayers of naive C6/36 cells in L-15 medium containing 1% heat-inactivated FBS, 10%TPB and 1.2% antibiotic (Penicillin G and Streptomycin). At days 5-7 after challenge, the supernatant solution was removed and used as inoculum for subsequent trials.

### 3.4 Acute and persistent infections of DENV-2 in C6/36

Persistent infections of DENV-2 in C6/36 cells were established as previously described ^5^. Briefly, we started with 10^6^ cells per well of naive C6/36 cells in 1 ml of culture media and incubated for 24 hours at 28ºC to achieve confluence. This was called passage zero (P0). The P0 were then challenged with DENV-2 at a multiplicity of infection (MOI) 0.1. After 2 hours of infection time, the medium was removed and replaced with fresh medium containing 2% FBS for further incubation at 28°C for 2 days. Then, all supernatant solutions were removed and cells were resuspended by knocking in fresh medium containing 10% FBS at 1:3 dilutions and transferred to new culture wells for 2-days cultivation /passage. This transfer constituted the 1^st^ passage. For subsequent passages, the decantation, suspension, dilution and transfer processes were carried out sequentially at 2-day intervals to establish persistently infected cultures. As previously reported ^5^, minor cytopathic effects (CPE) occurred for the first three passages (the acute infection stage) but were absent thereafter (the extended persistent infection stage). Three replicates were done in 6-well plates. Mock-infected cells were run in parallel to the viral infected cells to serve as negative controls.

### 3.5 MTT assay for TFD toxicity to C6/36 cells

Tenofovir [3-(4, 5-dimethylthiazol-2-yl)-2,5-diphenyl tetrazolium bromide] hereafter called TDF is as a nucleoside reverse transcriptase inhibitor in purified chemical powder form that was obtained from the Thailand Government Pharmaceutical Organization. Monolayers of C6/36 cells cultured for 24 hours were treated with TDF at concentrations of 0.1, 0.25 and 0.5 mM in culture media for 24 hours. The concentration used was calculated from the manufacturers recommended dose for human use. After 24 hours’ treatment with TDF, cells layers were examined and photographed with an inverted Olympus IX71 light microscope equipped with a digital camera that yielded photomicrographs with embedded scale bars. Then, culture media were removed and cell viability was tested using the MTT reduction assay ^10^. Briefly, cell monolayers treated with different concentrations of TDF were incubated with 100 μl MTT solution (0.5 mg/ml) for 2 hours at 28 °C protected from light. After incubation, the formazan salts formed were dissolved in dimethyl sulfoxide for 15 min followed by measuring intensity at 490 nm using a microplate reader (TECAN, Switzerland). Cultures of untreated cells were used as the negative control with their intensity considered as 100% relative viability.

### 3.6 Naive cell inhibition of reverse transcriptase (RT) upon DENV-2 challenge

Monolayers of C6/36 cells at Passage 0 were treated with TDF at different concentrations as described above for 24 hours. The treated monolayers were then challenged with DENV-2 at a multiplicity of infection (MOI) of 0.1. After incubation for 2 hours with gentle shaking at room temperature, the medium was removed, cells were washed 2 times with PBS and fresh medium containing 10% FBS was added for further incubation at 28°C for 48 hours (2 days) (acute infections in C6/36 cells). Then the supernatant solution was removed for DENV-2 copy number measurement, and cells were re-suspended and serially passaged as described above but with TDF in the medium at concentrations of 0.1, 0.25 and 0.5 mM. The negative control was untreated, unchallenged C6/36 cells and the positive control was untreated C6/36 cells challenged with DENV-2. There were 3 replicates for each group.

### 3.7 Inhibition of reverse transcriptase in cultures persistently infected with DENV-2

The persistently DENV-2 infected cultures were established by serial passage as described above. Monolayers of C6/ 36 cells persistently infected for 30 passages were treated with various concentrations of TDF for 48 hours. Then morphology of the whole culture was observed under the inverted light microscope (Olympus IX71) and DENV-2 copy number in the culture medium was measured in by real-time RT PCR.

### 3.8 One step real-time RT-PCR for DENV-2 quantification

Culture media or cells were collected for RNA extraction using an RNA extraction kit (Qiagen). DENV-2 copy number was quantified by one step real time RT-PCR as previously described ^5^. Briefly Primers D2S (5’-GTT CGT CTG CAA ACA CTC CA-3’) and D2C (5’-GTG TTA TTT TGA TTT CCT TG-3’) were used to obtain an expected amplicon of 230 bp and a standard curve of DENV-2 copy number was prepared by serial dilution of a plasmid containing the dengue virus envelope gene target. Each treatment was compared by gene copy number. One step real time RT-PCR was performed using 100 ng of total RNA and KAPA SYBR FAST One-Step qRT-PCR Kits (Kapa Biosystems) according to the directions provided by the manufacturer. Copy numbers were quantified per 100 ng total RNA template.

## Author contributions

Warachin Gangnonngiw: formal analysis, investigation, validation, writing – original draft and writing.

Timothy W Flegel: conceptualization, data curation, supervision, visualization and writing – review and editing.

Nipaporn Kanthong: conceptualization, data curation, methodology, supervision and writing – review and editing.

## Data availability statement

No data were generated during this research or is required for the work to be reproduced.

## Ethical statement

No human or animal research was conducted; thus, this study did not require approval from an Ethical Committee.

